# Taurine is a Natural Host Cytoadhesion Inhibitor in Asymptomatic Malaria Cases

**DOI:** 10.1101/2025.07.06.663375

**Authors:** Gretchen Diffendall, Perine Millot, Patty Chen, Balotin Fogang, Dominique Dorin-Semblat, Fanny Aprahamian, Sylvère Durand, Benoît Gamain, Antoine Claessens, Artur Scherf

## Abstract

The prolonged dry season in malaria-endemic regions of sub-Saharan Africa can be divided into periods of high and low transmission. The majority of symptomatic malaria cases are restricted to the short high transmission period that follows the rainy season. Shortly after, persistent asymptomatic malaria cases are more prevalent throughout the prolonged low transmission period. It is still unclear whether host metabolic alterations play a role in asymptomatic infections during seasonal malaria. In this study, we analyzed the blood plasma metabolome (n=199) of individuals in The Gambia, West Africa, capturing data from both high and low malaria transmission periods. Plasma samples from individuals (n=16) were collected monthly throughout the low transmission season, enabling a longitudinal analysis of metabolic alternations over six months. Our findings reveal that significant changes in host plasma metabolite composition are associated with seasonality and malaria pathogenicity. Notably, we observed elevated levels of taurine in asymptomatic malaria infections, especially during periods of low transmission. *In vitro*, this naturally occurring host molecule inhibits the cytoadhesion of malaria-infected red blood cells (iRBCs), which is key to malaria disease severity and mortality. Exogenous taurine can significantly reduce or reverse binding of iRBCs to the common adhesion receptor CD36 and the endothelial protein C receptor (EPCR), the later being associated with cerebral malaria. This study uncovers a mechanism by which elevated taurine plasma levels in asymptomatic infections could reduce cytoadhesion and lead to increased splenic clearance, thereby strengthening host resistance to symptomatic infections. In the absence of health strategies targeting dry season parasite reservoirs, our findings highlight taurine as a potential prophylactic or therapeutic agent to reduce symptomatic malaria in sub-Saharan Africa.

**One Sentence Summary:** Metabolomics uncovers taurine as a natural host inhibitor of *Plasmodium falciparum* cytoadhesion in asymptomatic malaria carriers, offering new insights for disease control.

## INTRODUCTION

Malaria, a mosquito-borne disease, remains a significant global health issue, causing nearly 600,000 deaths annually, with the majority occurring among children in sub-Saharan Africa ^1^. Most fatalities are due to infection by *Plasmodium falciparum*, which produces adhesive proteins known as PfEMP1 on the surface of infected red blood cells (iRBCs) ^2^. These proteins enable the parasite to bind to endothelial cells in various organs in a process called cytoadhesion, helping it to evade clearance by the spleen. The accumulation of parasites in organs is associated with symptomatic malaria, and when this occurs in the brain, it can lead to cerebral malaria, a more severe form of the disease ^3^. Due to its central role in virulence, PfEMP1 is one of the most extensively studied surface protein families, consisting of 60 members per genome. By switching expression between these family members, iRBCs can exhibit different adhesive properties and antigenic variants, enabling them to evade the immune system and to sequester in different organs ^4,5^.

The extracellular domain of PfEMP1 proteins is comprised of 2-10 Duffy-binding-like (DBL) domains and cysteine-rich interdomain regions (CIDR) ^6–9^. Individual domains commonly act as specific receptor-binding regions of endothelial surface receptors ^10^. Despite the significant sequence diversity among PfEMP1 family members, the N-terminal CIDRα domains retain a structural configuration that allows binding to either CD36 (CIDRα2-6) ^11,12^ or EPCR (CIDRα1) ^10,13,14^. CD36 is predicted to be the most common adhesive phenotype in the PfEMP1 family, since approximately 84% of PfEMP1 proteins bind CD36 ^10^. This scavenger receptor, commonly expressed on endothelial and innate immune cells, interacts with iRBCs from laboratory strains and field isolates. Apart from preventing splenic clearance, binding to CD36 expressed on dendritic cells may also inhibit the host innate immune response to infection ^15,16^. A subset of 5-7 PfEMP1 members are predicted to bind to EPCR on brain endothelial cells ^17,18^ and has been associated with severe childhood malaria ^13,19^ and notably cerebral malaria ^20,21^.

Several studies have demonstrated that environmental sensing is a crucial aspect of the pathogen, modulating its proliferation, virulence gene expression, and developmental pathways ^22–27^. In the context of environmental sensing, a phenomenon known as seasonal malaria in Sub-Saharan countries is of particular interest ^28,29^. In these malaria-endemic regions, asymptomatic *P. falciparum* infections during periods of low transmission alternate with symptomatic infections during periods of high transmission. The influence of environmental changes on parasite virulence regulation and clinical manifestations remains a compelling area of investigation. Several epidemiological studies investigating factors that decrease symptomatic infections have been performed in Mali. The antibody response in asymptomatic individuals in Mali did not provide evidence that the immune response plays a major role in asymptomatic infections ^29^. However, the same authors observed an extended blood circulation time of infected red blood cells (iRBCs) in asymptomatic individuals, suggesting that enhanced clearance by the spleen reduces parasite load and clinical symptoms. The factors contributing to this phenotype remain unknown ^29–32^.

Over the course of a year, The Gambia can be divided into periods of high transmission (late September through mid-December), when mosquitoes are more prevalent to transmit *P. falciparum* parasites to new human hosts, and periods of low transmission (late December through mid-September), when transmission is reduced (Fig. 1A) ^33^. Both high and low transmission periods occur during the dry season (mid-September through mid-June) whereas the wet season is limited to late June through early September. After the high rain levels cease in September, symptomatic infections are limited to the high transmission months. A sizeable fraction of infections remain chronic, resulting in asymptomatic infections being detected throughout the year, not limited to a specific period ^34^.

**Fig. 1:**
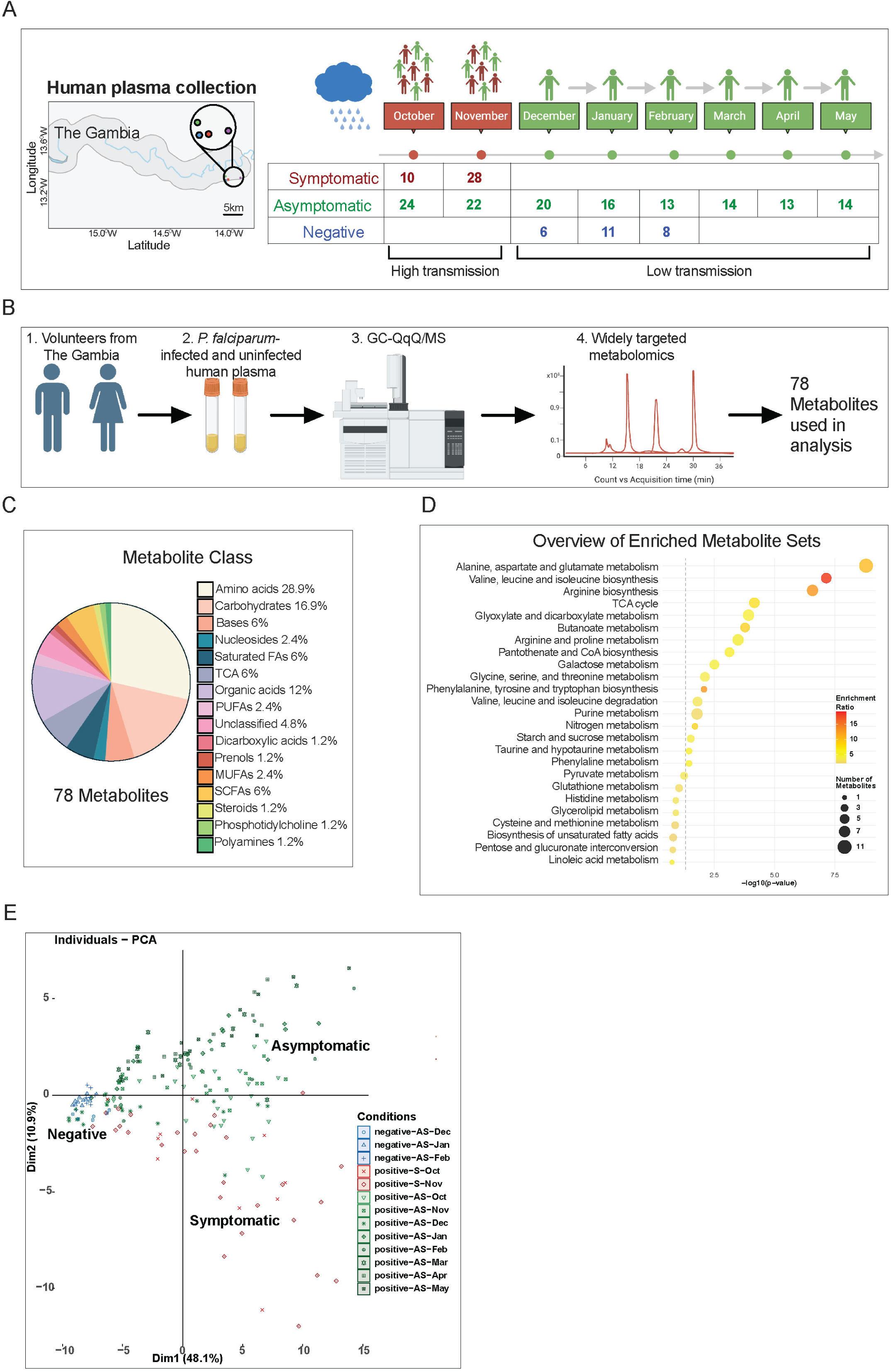
Blood Plasma Metabolomic Profiling of *P. falciparum* Infections in The Gambia. (**A**) Region in The Gambia where plasma samples were taken from volunteers in neighboring villages within a 5km radius (left). Overview of the change from high transmission to low transmission months in The Gambia. Sample numbers are indicated by each month for volunteers that had symptomatic or asymptomatic infections as well individuals with no detected presence of *P. falciparum* (right). (**B**) Schematic showing the flow of the experimental analysis for gas chromatography coupled with triple-quadrupole mass spectrometry (GC– QqQ/MS) for widely targeted metabolites. (A, B) Figures created with BioRender.com (**C**) Metabolite class enrichment for all 78 metabolites that passed quality control. (**D**) Pathway enrichment analysis of the 78 metabolites used in analysis are shown with p-value and enrichment ratio. (**E**) PCA plot for all 78 metabolites and all (n=199) plasma samples for the three clinical conditions (symptomatic infections, asymptomatic infections, and uninfected negative).

Although metabolomic analyses of human plasma from *P. falciparum*-infected individuals, have been conducted, most have focused on the effects of disease severity ^35,35–39^ rather than on seasonal variations in malaria. To our knowledge, only one recent study ^32^ has examined the impact of persistent asymptomatic infections on host plasma metabolites in Mali. However, its small sample size may have limited the ability to detect significant metabolic alterations.

In this study, we explored whether the dry season affects the host plasma environment of infected individuals, with a particular focus on identifying metabolites associated with disease severity and seasonal impact on asymptomatic infections. Metabolite profiling was performed on human plasma samples (n = 199) collected from individuals in The Gambia, including those infected with *P. falciparum*—both symptomatic and asymptomatic—as well as non-infected volunteers, as part of a previously conducted two-part study^33,40^.

Our results reveal significant changes in metabolite levels associated with malaria severity— comparing symptomatic and asymptomatic infections during the high transmission season—as well as with seasonality, by comparing asymptomatic infections across high and low transmission periods. Notably, one of the most highly elevated metabolites in asymptomatic infections, consistently increased in asymptomatic infections, was taurine, an amino sulfonic acid. While taurine is synthesized in the liver from methionine and cysteine, it is not a component of proteins but plays a role in various physiological host functions [reviewed in ^41^]. Although taurine showed no effect on iRBCs development of cultured parasites, it strongly inhibited iRBCs adhesion to CD36 and EPCR receptors as well as by a human cell line expressing CD36 at concentrations observed in dry season-infected individuals. Additionally, taurine was able to reverse iRBCs binding to these receptors. The identification of a natural host molecule that inhibits iRBC binding to CD36 and EPCR offers a novel therapeutic and prophylactic strategy to target symptomatic malaria and reduce dry-season reservoirs.

## RESULTS

### Study design and metabolomics analysis

To investigate host metabolic changes during malaria infection, we analyzed a total of 199 plasma samples collected from individuals living in neighboring villages in The Gambia (Fig. 1A) as part of a previously published longitudinal study ^33,40^. The cohort design included two complementary arms: (1) During the high transmission season (October–November), active case detection yielded 46 blood samples from asymptomatic *P. falciparum* carriers, while individuals presenting at local health posts yielded 38 samples from symptomatic malaria case. (2) During the low transmission season (December–May), a subset of 16 asymptomatic individuals with persistent *P. falciparum* infections was followed monthly for up to 6 months. Also included were 9 unique asymptomatic individuals that had samples taken in December. Additionally, 11 non-infected individuals were sampled between December to February.

As expected, parasite loads were significantly higher in symptomatic individuals (Fig. S1A). Although participants were not age-matched across groups, the overall age distribution was representative of the local population (median age: 19 years, Fig. S1B–C, ^33^). Metabolomics was performed on a total 199 plasma samples using gas chromatography coupled to triple quadrupole mass spectrometry (GC–QqQ/MS), targeting a broad panel of compounds. After quality filtering, 78 metabolites were retained for analysis (Fig. 1B), spanning diverse chemical classes including amino acids, fatty acids, and carbohydrates (Fig. 1C). These metabolites play a role in various metabolic pathways, determined by pathway enrichment analysis (Fig. 1D, SM3). Initial Principal component analysis (PCA) revealed clustering of samples according to infection status and season (Fig. 1E).

### Metabolites associated with seasonality

We assessed the influence of seasonality on host metabolic responses by comparing asymptomatic infections from the peak transmission period (October) with those from the end of the dry season (May), when rainfall has been absent for over seven months and malaria transmission is nearly absent. Seventeen metabolites differed significantly between these timepoints (Wilcoxon test, adjusted for multiple comparisons with Benjamini-Hochberg correction, Fig 2A, SM5). The most significant increase in May was observed for taurine (adjusted p-value < 0.00379). These results indicate that the dry season influences host plasma metabolites, possibly through changes in diet or microbiota composition.

**Fig. 2:**
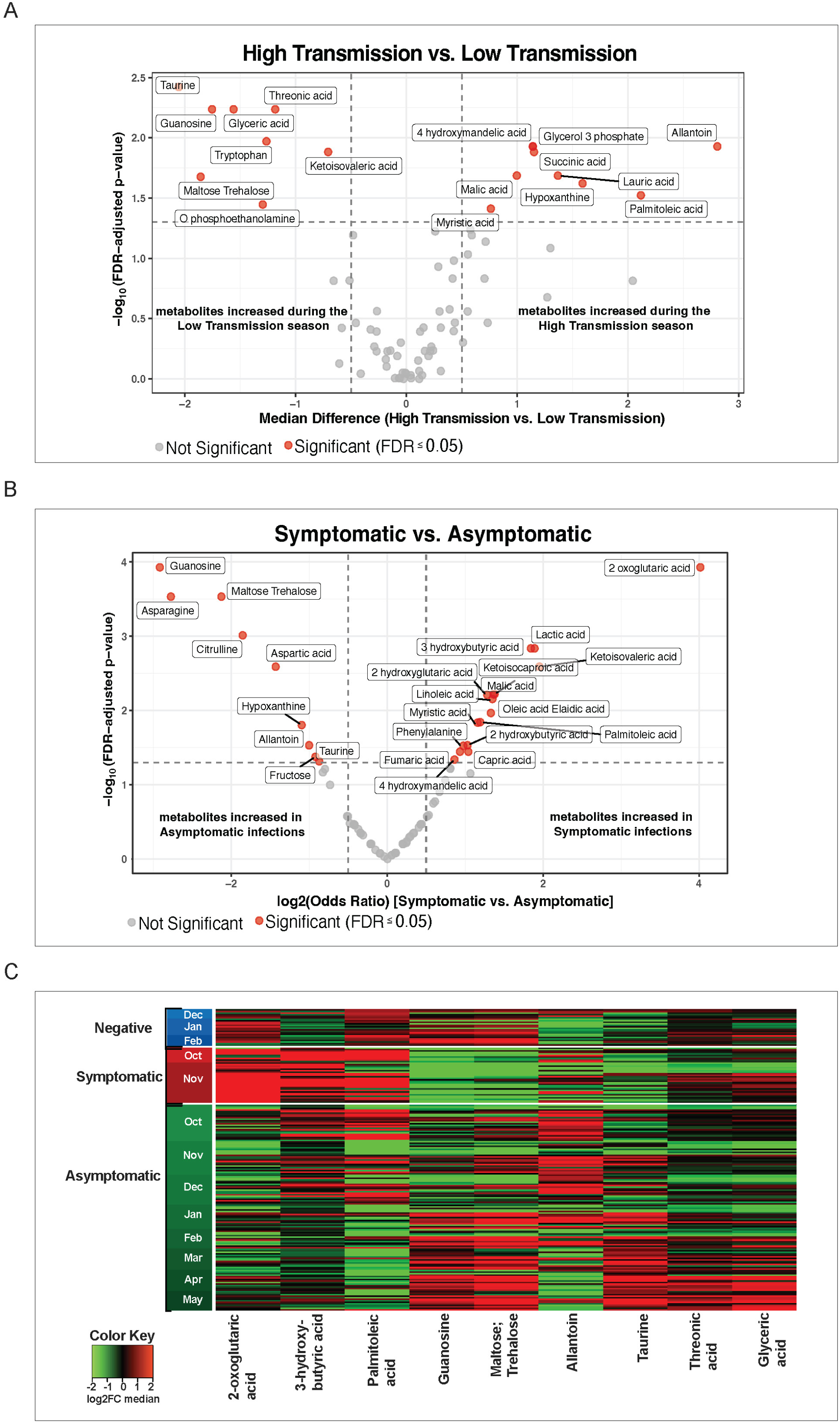
Metabolites associated with seasonality and severity. (**A**) Volcano plot showing metabolites that had significantly increased levels in asymptomatic infections during the high transmission season (October) compared to the low transmission season (May). The median difference is plotted on the x-axis and the -log_10_ (FDR adjusted p-value) is plotted on the y-axis. Red colored circles represent significant metabolites with FDR ≤ 0.05 **(B)** Volcano plot showing metabolites that had significantly increased levels in symptomatic infections during the high transmission season (October, November) compared to asymptomatic infection during the high transmission season (October, November). The log_2_(Odds ratio) is plotted on the x-axis and the -log_10_ (FDR adjusted p-value) is plotted on the y-axis. Red colored circles represent significant metabolites with FDR ≤ 0.05 **(C)** Heatmap from MS analysis showing metabolites that varied based on severity and seasonality. All (n=199) samples, months, and conditions are included. Columns are metabolites, rows are samples. Heatmap data are log2 normalized (within a range of [-2]-[2]) and centered around the average abundance computed from all the samples for each metabolite. Red/green colors are ion signal higher/lower than average. Samples and metabolites are clustered following the ward.D2 algorithm, with Euclidean distance. **(C)**

### Metabolites associated with symptomatic malaria

To identify metabolic signatures associated with malaria symptoms, we compared 38 individuals with symptomatic infections to 46 asymptomatic infections collected during the same high-transmission months (October–November). We used a mixed-effects model with month as a random effect and metabolites as fixed effects. After multiple testing correction, 27 metabolites were significantly associated with clinical status, with 16 elevated and 11 reduced in symptomatic individuals (Fig. 2 B&C, SM5). Among the metabolites elevated in symptomatic individuals were: (1) 2-oxoglutaric acid (more commonly known as alpha-ketoglutarate, a TCA cycle intermediate, previously identified as a biomarker of *P. falciparum* infection ^36^. (2) lactic acid, consistently reported as elevated in symptomatic malaria. (2) Palmitoleic acid, a monounsaturated fatty acid that may accumulate during *P. falciparum* infection ^42^. (4) 3-hydroxybutyric acid, a fatty acid derivative that accumulates during parasite-induced acidosis ^43^ and has been shown to inhibit *P. falciparum* growth *in vitro* ^44^.

Conversely, multiple amino acids (aspargine, citrulline, aspartic acid) and other metabolites (guanosine, maltose;trehalose) were reduced in symptomatic infections. Taurine levels were also lower, though less markedly (Odds Ratio 0.553, adjusted p-value = 0.0503). These findings reveal a distinct metabolic host signature associated with symptomatic malaria, highlighting disruptions in energy metabolism and fatty acid pathways during acute *P. falciparum* infection.

### Metabolite dynamics during chronic asymptomatic infection in the dry season

We took advantage of the longitudinal monthly sampling from 16 individuals with persistent asymptomatic infections over the low transmission season (December to May) to examine temporal changes in plasma metabolites. Using a mixed-effects model with non-infected individuals as controls, we identified nine metabolites—including several amino acids and taurine—that consistently increased over time, particularly from March onward (Fig S2A, SM5). In contrast, allantoin and glycerol-3-phophate showed the opposite trend, with higher levels early in the low transmission season and lower levels later (Fig. S2A, SM5). These patterns likely reflect a combination of dietary shifts and host physiological adaptations. For instance, allantoin, an oxidative product of urate, is a marker of oxidative stress; valine, isoleucine and methionine are diet-derived; taurine can be either diet-derived or endogenously synthesized, and plays a protective role against oxidative stress ^45^.

Finally, we used a mixed-effects model to assess whether the host metabolome differentiates asymptomatic carriers during the low transmission season from uninfected individuals. Only two metabolites, capirc acid and aminovaleric acid, were associated with infection status, indicating a weak effect of an asymptomatic infection on the overall metabolome (SM5). Nevertheless, the dynamic shift in specific plasma metabolites across the low transmission season likely reflect dietary changes and host physiological adaptations, including responses to oxidative stress.

This dataset motivated us to identify metabolites potentially involved in modulating *P. falciparum* persistence and virulence. We prioritized metabolites meeting three progressively less stringent criteria: (1) higher levels in May compared to the peak transmission season (October), (2) a consistent increase across the low transmission season, (3) lower levels in asymptomatic than in symptomatic cases. Taurine emerged as the strongest candidate, fulfilling all three conditions (Fig. S2B). These findings led us to further investigate the functional role of taurine in malaria pathogenesis, a direction supported by a recent study showing that taurine supplementation in a *P. berghei* mouse model conferred host benefits through immune modulation ^46^.

### Taurine inhibits ability of iRBCs to cytoadhere to CD36 and EPCR

Given the key role of iRBC adhesion in malaria pathogenesis, our goal was to identify candidate metabolites significantly elevated in asymptomatic infections during the low transmission period that may influence *P. falciparum* sequestration to endothelial receptors. Additionally to taurine, threonic acid, a breakdown product of ascorbic acid, was selected due to previous findings that ascorbic acid significantly inhibits *P. falciparum* growth; however, its potential effect on iRBC cytoadhesion has not yet been studied ^47^. We next investigated whether taurine and threonic acid could reduce iRBC cytoadhesion *in vitro*. To establish physiologically relevant high and low taurine concentrations for the cultures, we measured the absolute taurine quantity in plasma samples exhibiting the highest and lowest values from our mass spectrometry analysis. The lowest values corresponded to taurine concentrations of [0.009mM], [0.026mM], and [0.032mM] while the highest values corresponded to [0.113mM], [0.175mM], and [0.176mM]. These taurine concentrations aligned with levels reported in human plasma ^48^ and confirmed the mass spectrometry results. Due to the absence of an assay kit for threonic acid, a reported physiological concentration of 0.27 mM ^49^ was used for all analyses. Growth curve analysis of wild-type *P. falciparum* cultures supplemented with or without various taurine concentrations, or threonic acid [0.27mM] (Fig. S3A) revealed no effect of taurine nor threonic acid on growth over the course of five days. These data are consistent with what was observed in another study ^50^ at even higher concentrations of taurine. Physiological taurine levels ranging from 0.64 to 0.93 mM have been observed in clinical volunteer studies following oral administration of 4 g of taurine ^51^.

To determine if adhesion of iRBCs is affected by taurine or threonic acid, we performed adhesion assays with RBCs infected with late-stage wild-type 3D7 parasites grown in media supplemented with a range of taurine concentrations for up to 40 hours (Fig. 3A, S3B). The lowest taurine concentration of [0.02mM] already significantly decreased adhesion of iRBCs to CD36 (p < 0.0001) by up to 60% (Fig. 3A). With even higher taurine concentrations, binding was drastically decreased by > 97% (Fig. 3A, B). Thus, physiological levels of plasma taurine found in asymptomatic individuals almost completely blocks binding of iRBCs to recombinant CD36 receptor *in vitro*. Threonic acid, at physiological levels of [0.27mM] ^49^, had no significant effect on adhesion of iRBCs to CD36 (Fig. S3C). As metabolomic MS analysis revealed no significant differences in methionine levels between conditions (SM4), a physiological concentration of 0.04 mM was used as a control, which also had no significant effect on adhesion (Fig. 3A) ^52^. Incubation of iRBCs with [0.16mM] of taurine for one hour prior to the adhesion assays was sufficient to decrease cytoadhesion by > 97% (Fig. 3A), suggesting that taurine influences iRBC adhesion through direct interaction with the iRBC surface, rather than by altering parasite transcription or protein expression on the surface of iRBC.

**Fig. 3:**
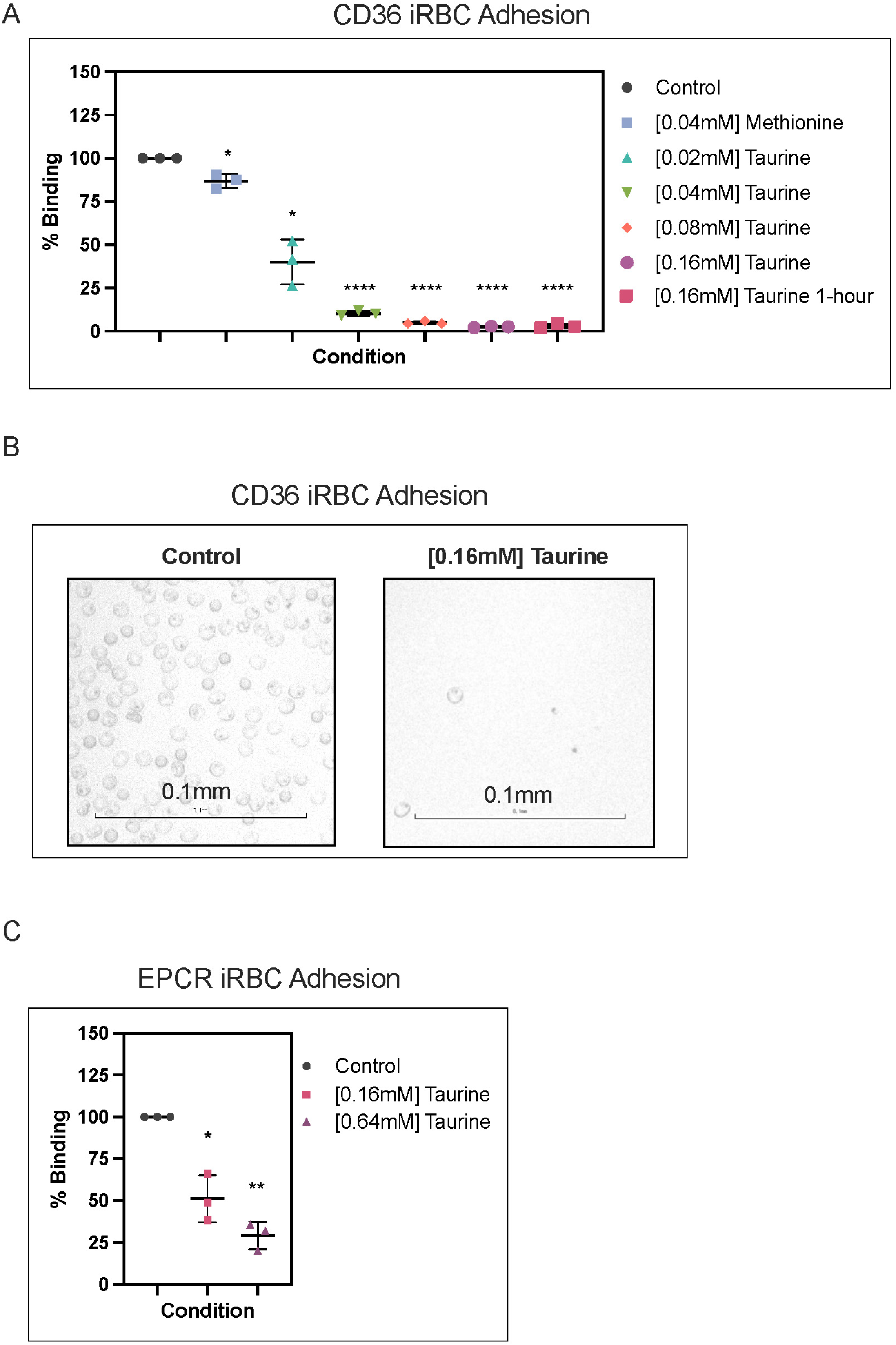
Taurine markedly reduces iRBC adhesion. (**A**) iRBC adhesion binding assays showing 3D7 iRBCs binding to CD36. Parasites have been incubated for 30 hours in [0.04mM] methionine or increasing concentrations of taurine. Additionally, parasites were incubated for one hour in [0.16mM] taurine. Data is expressed as the percentage of iRBCs binding ±SD per mm^2^ for each condition compared to control, with the median represented at the center line. Results shown are three biological replicates (n=3) from duplicate technical replicates. Statistical significance was determined by paired t-test (* p = 0.0305 and 0.0152, **** p < 0.0001). (**B**) Representative iRBC adhesion assay images are shown for control and supplementation of taurine physiological concentration [0.16mM]. (**C**) EPCR binding assay of Itvar19 FCR3 iRBCs in the presence of [0.16mM] taurine and [0.64mM] taurine. Data is expressed as the percentage of iRBCs binding ±SD per mm^2^ for each condition compared to control, with the median represented at the center line. Results are from a total of three biological replicates (n=3) done in duplicate. Statistical significance was determined by paired t-test (* p = 0.0264, ** p = 0.0045).

The surface PfEMP1 domains that bind to CD36 and EPCR are associated with a single CIDR domain ^53^. We next set out to determine if taurine affects iRBC binding to the endothelial cell protein C receptor (EPCR), which is important in cerebral malaria ^13^. We used a parasite strain (FCR3) that predominantly express the Itvar19 PfEMP1 previously found to bind to EPCR ^54^. After a one-hour incubation with [0.16mM] or [0.64mM] taurine, we observed a significant decrease in adhesion of FCR3 iRBCs to recombinant EPCR of almost 50% and >70%, respectively (Fig. 3C). This parasite line, which also expresses a distinct PfEMP1 member was additionally used for adhesion assays with CD36 (Fig. S3D). Taurine supplementation similarly reduced receptor binding across strains, indicating that taurine inhibitory effect is not strain specific and is effective despite differences in *var* gene family repertoires.

In contrast to adhesion of infected red blood cells to CD36 and EPCR, which involves diverse PfEMP1 variants containing CIDR domains, adhesion to placental chondroitin sulfate A (CSA) is mediated by a unique PfEMP1 encoded by the *var2CSA* gene, which comprises multiple DBL domains but lacks a CIDR domain ^53^. We performed iRBC adhesion assays with the NF54 parasite strain that expresses VAR2CSA ^55^ in the presence or not of [0.64mM] taurine (Fig. S3E). A small but significant decrease in iRBC binding was observed (p = 0.0288). Finally we assessed the capacity of malaria naïve human serum to inhibit iRBC binding to CD36. Human serum caused a slight reduction in our iRBC binding assay (Fig. S3F), comparable to the effect of 0.02 mM taurine supplementation (Fig. 3B), likely due to the taurine content in the serum used for parasite culture, which reached up to 0.02 mM. Our findings show that physiological taurine levels inhibit iRBC adhesion to CD36 and EPCR, pointing to a novel host-mediated mechanism contributing to reduced *P. falciparum* virulence in asymptomatic infections.

### Taurine reverses adhesion of iRBCs to CD36 and EPCR

To assess its therapeutic potential, we set out to determine if taurine could reverse, rather than merely prevent, iRBC binding to CD36 (Fig. 4A) and EPCR (Fig. 4B), using the same strains from (Fig. 3B, D). When iRBCs already adhering to CD36 or EPCR were incubated without or with [0.64mM] taurine for 45 minutes, adhesion decreased by over 77% (p < 0.0001) to CD36 and nearly 30% (p = 0.0017) to EPCR (Fig. 4A, B). These demonstrate that taurine can disrupt existing interactions of iRBCs with endothelial receptors.

**Fig. 4:**
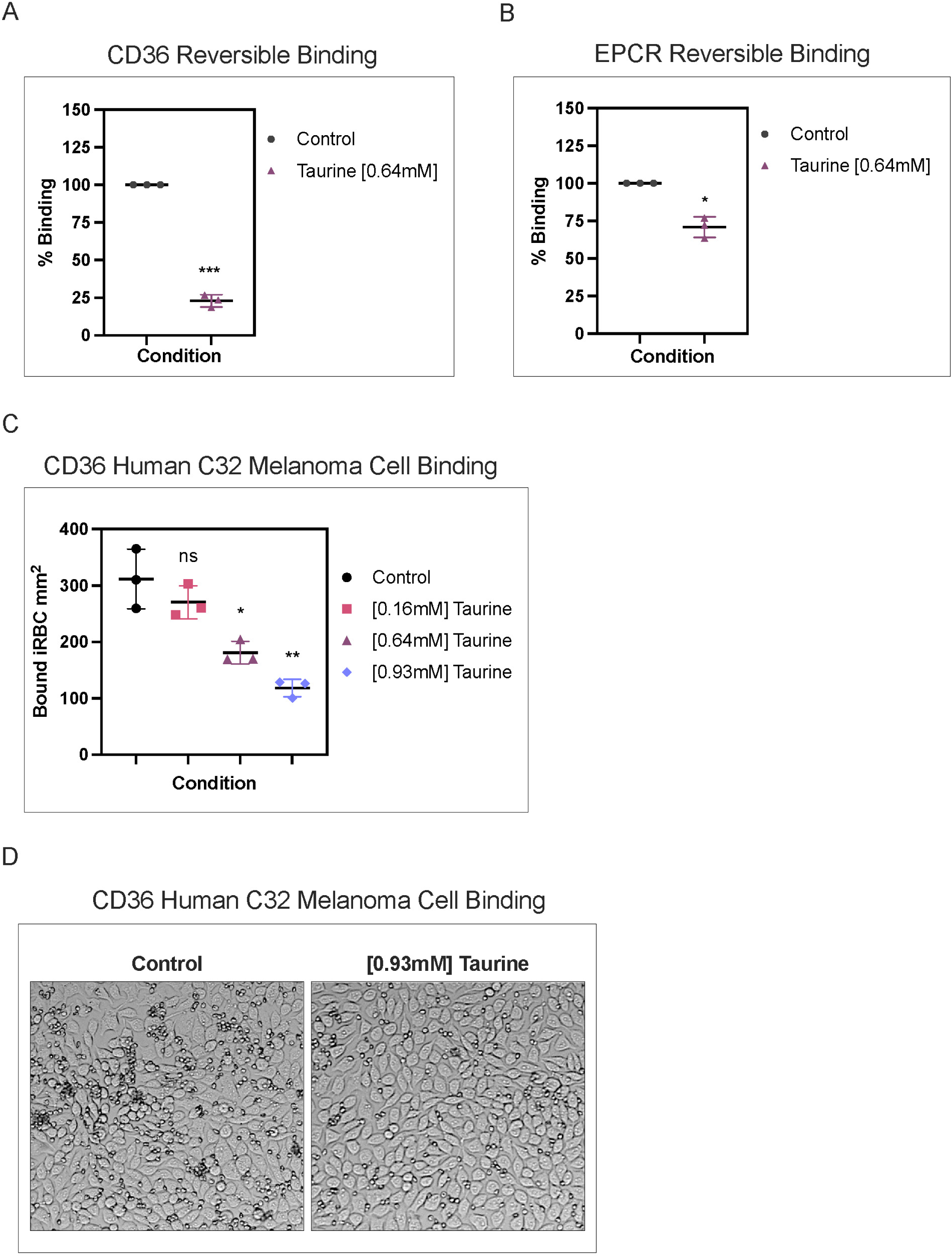
Taurine-Mediated Reversal of iRBC Receptor Binding and Inhibition of Cytoadhesion to C32 Cells. (**A**) Reversal of iRBC adhesion of 3D7 iRBCs bound to CD36, incubated with or without [0.64mM] taurine. Data is expressed as the fold change of bound iRBCs ±SD per mm^2^ compared to control, with the median represented at the center line. Results are from a total of three biological replicates (n=3) done in duplicate. Statistical significance was determined by paired t-test (*** p = 0.0009). (**B**) Reversible iRBC adhesion assays for FCR3 Itvar19 parasites bound to EPCR incubated with or without [0.64mM] taurine. Data is expressed as the fold change of bound iRBCs ±SD per mm^2^ compared to control, with the median represented at the center line. Results are from a total of three biological replicates (n=3) done in duplicate. Statistical significance was determined by paired t-test (** p = 0.0179). (**C**) Cytoadhesion assays measuring *P. falciparum* FCR3 parasite strain binding to human C32 melanoma cells expressing CD36 and ICAM-1 under control conditions and with increasing taurine concentrations (0.16 mM, 0.64 mM, 0.93 mM). Data is expressed as the fold change of bound iRBCs per mm² of confluent C32 cells (±SD), with medians indicated by center lines. Results represent three biological replicates (n = 3), each in duplicate. Statistical significance was determined by paired t-test (* p = 0.0231, ** p = 0.0076); NS = not significant. (**D**) Representative field of cytoadhesion assay from (Fig. 4C).

To explore the mechanisms by which taurine inhibits and reverses iRBC binding to endothelial receptors, we examined PfEMP1 expression. Since a one-hour taurine incubation was sufficient to nearly abolish completely iRBC adhesion, it is unlikely that transcriptional changes could account for such rapid effects on protein expression. Indeed, FACS analysis of FCR3 Itvar19 iRBCs supplemented with taurine showed no significant changes in surface PfEMP1 protein levels (Fig. S4A, S4B, C). We additionally tested whether taurine interacts directly with CD36 to prevent iRBC binding. iRBC adhesion assays were performed with CD36 receptors that were pre-treated with or without [0.16mM] taurine and no significant difference in adhesion of iRBCs was observed between the two conditions (Fig. S4D). The observed rapid action of taurine suggests it may directly target the iRBC surface, inducing conformational changes in parasite adhesins and thereby altering binding properties.

### Taurine reduces binding of iRBCs to CD36 on human C32 melanoma cells

To investigate taurine inhibition in a cellular context, we employed the human C32 amelanotic melanoma cell line, which has been used extensively in previous *P. falciparum* CD36 adhesion studies ^56^. This cell line shares adhesion receptors with human endothelial cells, and the mechanisms of binding are comparable, making it a suitable surrogate for endothelial cells in adhesion assays ^57,58^. Parasitized erythrocytes primarily adhere to these melanoma cells via the glycoprotein CD36, which is also widely expressed on vascular endothelium ^59^. In three biological replicates we observed > 60% inhibition of FCR3 strain adhesion to C32 at taurine concentration of 0.93 mM. This inhibition was dose-dependent (Fig. 4C, D). Similar C32 inhibition results were obtained using a genetically distinct strain (3D7) (Fig. S4E, F). Taurine inhibits iRBC adhesion at the cellular level, albeit at higher concentrations than with recombinant receptors. Its effectiveness in reducing iRBC binding by multiple methods, highlights its potential as a natural supplement to lessen disease severity in endemic areas.

## DISCUSSION

The mechanisms by which *P. falciparum* persists through the rainy season without causing symptoms remain unclear. In this study, we investigated if environmental factors contribute to asymptomatic and persistent *P. falciparum* infections during the dry season in sub-Saharan Africa. To do so, we conducted a metabolomics analysis on plasma samples collected from infected and non-infected individuals in The Gambia, West Africa, from both high and low transmission times during the dry season (Fig. 1). Of the statistically validated metabolites identified (n=78), forty-three showed distinct profiles in the cohort (Fig. 2C). We identified metabolites that are dependent on disease severity or seasonality. Metabolites associated with severity include fatty acids, which have been shown to increase in *P. falciparum* infections, as well as a previously identified biomarker involved in the TCA cycle ^36^. Metabolites linked to seasonality include those potentially derived from the diet, as well as metabolites produced endogenously by the host to combat oxidative stress.

Since the development and proliferation of *P. falciparum* rely on nutrient uptake from the host serum ^60^, the observed changes in the host metabolome may directly affect the parasite. However, it remains unclear whether the altered metabolite levels—either elevated or reduced—are taken up by iRBCs and subsequently influence parasite development. Of note, a recent report suggests that external lactate levels, a major cause of parasite-induced acidosis, may serve as metabolic sensors in *P. falciparum* iRBCs ^61^. An increase in environmental lactate was linked to elevated levels of epigenetic regulators, *sirtuins* ^62^, and histone lysine lactylation ^61^ in the parasite. In our study, lactate levels were significantly lower in asymptomatic infections compared to those with symptomatic infections during the high transmission period. Another major contributor to parasite-induced acidosis, 3-hydroxybutyric acid, was found significantly increased in our symptomatic high transmission infections ^43^.

One of the most striking metabolic changes observed was in taurine levels, which were as low as 0.02 mM during periods of high transmission and rose to 0.16 mM by the end of the low transmission season in infected individuals. For comparison, oral administration of taurine (4 g) in eight healthy volunteers increased plasma levels from 0.03–0.06 mM to peak concentrations of 0.53–0.93 mM for several hours after taurine administration ^51^. The cause of elevated taurine levels in individuals with asymptomatic infections remains unclear. Although the human liver synthesizes taurine, dietary sources such as meat, fish, poultry, and dairy also contribute to plasma levels. In rural Gambia, diets mainly consist of rice, wheat-flour bread, and groundnuts, with seasonal availability of leafy vegetables, millet, and dried fish. Since there’s no evidence of increased protein intake during the dry season in this cohort, we hypothesize that elevated plasma taurine levels are due to increased endogenous bile secretion. Notably, persistent high host-derived taurine levels have been reported for extended periods following the resolution of enteric infections in mice ^63^. It would be important to investigate whether changes in the gut microbiome play a role in stimulating increased bile taurine production in this cohort.

Although taurine treatment did not affect parasite growth (Fig. S3A), it strongly inhibited the binding of iRBCs to recombinant receptor CD36 (Fig. 3A) and to a lesser degree to the brain endothelial receptor EPCR (Fig. 3C). Conversely, taurine had a very mild effect on adhesion of RBCs infected with parasites expressing an atypical PfEMP1 family member (VAR2CSA) that has no CIDRα domain (Fig. S3E). Importantly, we show that taurine inhibits iRBC adhesion to the human C32 melanoma cell line, a common endothelial surrogate in *P. falciparum* CD36 adhesion assays (Fig. 4C).

Since taurine preincubation with CD36 did not affect iRBC binding (Fig. S4D), it likely disrupts adhesion by targeting PfEMP1–receptor interactions, possibly at the CIDR-binding domain. Multiple findings support this hypothesis. Taurine pretreatment of adhesive iRBCs for one hour significantly reduced CD36 binding (Fig. 3A) without affecting PfEMP1 surface expression (Fig. S4A), suggesting the effect is not due to altered protein trafficking. Its selective inhibition of CD36 and EPCR binding points to disruption of specific PfEMP1 interactions rather than general membrane effects. Notably, taurine reversed adhesion to CD36 by over 77% and to EPCR by over 29% (Fig. 4A, B). For CD36, binding inhibition was consistent across CD36-binding strains (3D7 and FCR3), despite sequence variability in the CIDRα domain ^11^.

A study conducted in Mali^29^ provided evidence that increased circulation time of infected red blood cells (iRBCs) is associated with asymptomatic infections during the dry season, although the underlying mechanism was not elucidated. Our findings from The Gambia offer mechanistic support for these observations, suggesting that elevated plasma taurine levels in persistent asymptomatic infections may reduce or reverse iRBC binding, thereby promoting the circulation of mature parasite forms and their clearance by the spleen. In a recent small-scale plasma metabolomics study involving 35 samples from 20 individuals with symptomatic and asymptomatic infections in Mali, taurine levels were not reported ^32^. Further studies are needed to explore whether elevated plasma taurine levels are a consistent feature of asymptomatic *P. falciparum* infections during the dry season across sub-Saharan Africa. It cannot be excluded that, in addition to taurine, other metabolites associated with plasma alterations in asymptomatic malaria contribute to asymptomatic malaria by influencing the iRBC proliferation or by increasing their susceptibility to immune-mediated clearance.

Other host receptors such as ICAM-1 have been linked to severe malaria and aggregation of iRBCs and uninfected RBCs (known as rosettes) [reviewed in ^2^]. Both adhesion phenotypes are significantly rarer than CD36 adhesion and it remains to be shown if these phenotypes play an important role in symptomatic infections during seasonal malaria.

Binding of *P. falciparum* iRBCs to human receptors is central to the development of symptomatic and severe malaria. Yet, no adjunctive therapies or broad-spectrum vaccines currently target CD36-binding parasites, and efforts to block EPCR-binding variants implicated in severe and cerebral malaria remain ongoing ^64^. Clinically, taurine is already used as a dietary supplement with documented benefits for conditions like hypertension and hypercholesterolemia [reviewed in ^65^]. Our identification of taurine, a natural host molecule, as an effective inhibitor of iRBC binding to both CD36 and EPCR reveals a novel public health strategy to reduce or eliminate dry-season reservoirs and prevent symptomatic malaria at the start of the transmission season.

## MATERIALS AND METHODS

### Study individuals and ethical approval

Study design and sample collection was alredy reported in ^33,34,40^. The first part of the study was a cross-sectional study implemented in October and November 2016 to estimate the prevalence of *P. falciparum* infections in four neighboring villages (Madina Samako: K, Njayel: J, Sendebu: P, and Karandaba: N) within a 5km radius in Upper River Region (URR), eastern Gambia. *P. falciparum* asymptomatic infections were detected by varATS qPCR and individuals were invited to provide a venous blood sample before receiving anti-malarial treatment. Symptomatic cases presented themselves at the local health facility with fever. Malaria cases were defined as anxially temperature >37.5C, positive RDT and later confirmation of *P. falciparum* infection by varATS qPCR. For the second part of the study, in December 2016, asymptomatic *P. falciparum* carriers were recruited for a longitudinal cohort with monthly samplings until May 2017, or until the infection naturally cleared [Ref Collins2022].

The high transmission season months include October and November and the low transmission season months include late December, January, February, March, April, and May. The months pertaining to the two transmission periods was directly determined by the reports of symptomatic malaria infections for that specific year for all villages used in the study ^33^. For additional information about each sample, see (SM1).

The study protocol was reviewed and approved by the Gambia Government/MRC Joint Ethics Committee (SCC 1476, SCC 1318, L2015.50) and by the London School of Hygiene & Tropical Medicine ethics committee (Ref 10982). The field studies were also approved by local administrative representatives, the village chiefs. Written informed consent was obtained from participants over 18 years of age and from parents/guardians for participants under 18 years. Written assent was obtained from all individuals aged 12–17 years.

### Human plasma sample collection

From *P. falciparum* positive individuals (by varATS qPCR), 5 to 8mL venous blood was collected in a EDTA vacutainer. Collected venous blood samples were immediately separated in plasma, PBMCs and red blood cells, by centrifugation on a layer of Lymphoprep^TM^ according to manufacturer’s instructions. Plasma was directly stored at -80 degrees Celsius and never thawed until use in this study.

### Metabolomics of human plasma sample preparation and widely targeted detection by GC-MS

50 µl of collected plasma were mixed with 500 µl of ice-cold extraction mixture (methanol/water, 9/1, -20°C, with labelled internal standard). To facilitate endogenous metabolites extraction, samples were then completely homogenized (vortexed 5 minutes at 2500 rpm) and then centrifuged (15 min at 10000 g, 4°C). 150 µl of supernatants were collected to be analyzed by gas chromatography coupled with mass spectrometer (GC/MS) ^66^.

Widely targeted analysis by GC-MS/MS was performed on a 7890A gas chromatography (Agilent Technologies) coupled with Triple Quadrupole 7000C (Agilent Technologies) and was previously described in ^67^. All targeted treated data were merged and cleaned with a dedicated R (version 4.0) package (@Github/Kroemerlab/GRMeta).

### Plasma taurine concentration assay

Plasma taurine levels were determined using a commercial taurine assay kit (Cell Biolabs MET-5071). Complete information on tested individual samples can be found in (SM1). The protocol was followed as described in the kit and read on a Synergy 2 microplate reader for spectrophotometric reading.

### Parasite culture and synchronization

Asexual blood stage *P. falciparum* parasites were cultured as previously described in ^68^. Parasites were cultured in human RBCs (obtained from the Etablissement Français du Sang with approval number HS 2019-24803) in RPMI-1640 culture medium (Thermo Fisher 11875) supplemented with 10% v/v Albumax I (Thermo Fisher 11020), hypoxanthine (0.1 mM final concentration, C.C.Pro Z-41-M) and 10 mg gentamicin (Sigma G1397) at 4% hematocrit and under 5% O_2_, 3% CO_2_ at 37 °C. Static parasite development was monitored by Giemsa staining. Parasites were synchronized by sorbitol (5%, Sigma S6021) lysis at ring stage, plasmagel (Plasmion, Fresenius Kabi) enrichment of late stages 24 hours later, and an additional sorbitol lysis 3 hours after plasmagel enrichment. The 0-hour time point was considered to be 1.5 hours after plasmagel enrichment.

### Parasite growth assay

Parasite growth was measured as described previously ^69^. Wild type 3D7 parasites were tightly synchronized and diluted to 0.2% parasitemia (5% hematocrit) at the ring stage using the blood of three different donors separately (n=3). Each culture was split into control and parasites supplemented with [0.16mM] taurine. The growth curve was performed with three technical replicates per condition per blood. Parasitemia was measured every 24 hours by counting ten randomly selected different fields on Giemsa-stained slides each day for a total of five days.

### Static iRBC adhesion binding assays

<u>3D7-CD36 Binding Assay:</u> Mature-stage iRBCs (3D7 strain) were used for iRBC adhesion assays as previously described ^70^. Taurine was supplemented at increasing concentrations (0.02, 0.04, 0.08, and 0.16 mM) from a 200 mM stock to cultured parasites. Methionine (0.04 mM) was added post-synchronization. For short-term treatments, 0.16 mM taurine or 0.27 mM threonic acid was added one hour prior to the assay. All compounds were removed before adhesion assays via plasmagel enrichment of mature iRBCs. Recombinant receptors diluted in Dulbecco’s phosphate-buffered saline (DPBS, Thermo Fisher 14190) (10 µg/mL target CD36 or negative control 1% BSA) were spotted on labeled petri dishes overnight at 4°C in duplicate. Dishes were blocked for 30 minutes at 37°C with 1% BSA/DPBS while parasitemia was determined after trophozoite and young schizont iRBCs were isolated by plasmagel (Plasmion, Fresenius Kabi) enrichment. Adjusted amounts of iRBCs were resuspended in DPBS for a concentration of 2.2 × 10^8^ iRBCs/mL. iRBCs/DPBS were added to each petri dish and incubated for one hour at 37°C. Unbound cells were removed, and the dishes were washed five times with DPBS by carefully tilting from side to side. Adherent iRBCs were counted in duplicate spots using 40x lens with a Nikon ECLIPSE TE200 in seven randomly selected fields (each with 0.2 mm^2^) using ImageJ2 (version 2.14.0/1.54i). A total of three biological replicates were performed in duplicate and the results were expressed as the percentage of bound iRBCs compared to the control per 0.2 mm^2^ of target receptor monolayer.

#### FCR3-*var19*-EPCR and FCR3-CD36 binding assay

For EPCR binding, the protocol listed above was followed with the following changes: Recombinant receptors were diluted in in DPBS (50 µg/mL EPCR, 50 µg/mL CD36 or negative control 1% BSA) were spotted on labeled petri dishes overnight at 4°C in duplicate. Late stage iRBCs from FCR3 *var19*-expressing parasites ^54^ were purified by VARIOMACS (Miltenyi Biotec France) and counted. The iRBC pellet was split into control and addition of taurine [0.64mM] for one hour in RPMI-1640 culture medium (Thermo Fisher 11875) supplemented with 10% v/v Albumax I (Thermo Fisher 11020), hypoxanthine (0.1 mM final concentration, C.C.Pro Z-41-M) and 10 mg gentamicin (Sigma G1397) at 4% hematocrit and under 5% O_2_, 3% CO_2_ at 37 °C. The spots were washed twice with DPBS and blocked for 30 minutes at room temperature with 1% BSA/DPBS, 3.75 x 10^6^ parasites/mL diluted in 10uL DPBS were added to each spot and incubated for one hour at room temperature. Plates were gently washed with 25mL DPBS on an orbital shaker for 2 minutes and repeated four times before visualization and imaging using EVOS 5000. Adherent iRBCs were counted in duplicate spots using 20x lens with a Nikon ECLIPSE TE200 in seven randomly selected fields (each with 0.3 mm^2^).

#### NF54-*var2CSA*-CSA binding assay

For CSA binding, the protocol listed above (for 3D7-CD36 binding) was followed for control and supplementation of [0.64mM] Taurine using CSA-panned NF54 parasites ^55,71^.

#### Evaluation of taurine’s ability to reverse iRBC binding to CD36 and EPCR

A two-step protocol was used to evaluate the reversal of binding of 3D7 and FCR3-*var19*-expressing parasites to CD36 or EPCR. In Step 1, binding was conducted as previously described for 3D7 to CD36 and FCR3-*var19* to EPCR. In Step 2, DPBS or [0.64mM] taurine + DPBS was added for 45 minutes, followed by DPBS washes. Adherent iRBCs were counted in duplicate spots using a 10x objective lens to count and assess the percent reduction in bound iRBCs after taurine incubation compared to the control.

#### Taurine-CD36 incubation prior to iRBC adhesion assay

For CD36-Taurine binding, the protocol was followed the same as listed above (for 3D7-CD36 binding) for mature stage iRBCs from 3D7 parasites with the following changes: Control and supplementation of [0.16mM] taurine was added during blocking stage with 1% BSA/DPBS.

#### Human serum- and human serum+taurine-CD36 binding assay

For human serum-CD36 binding and human serum + taurine-CD36 binding, the protocol was followed the same as listed above (for 3D7-CD36 binding) for mature stage iRBCs from 3D7 parasites with the following changes: 10% human serum without and with supplementation of [0.64mM] taurine from three different donors (n=3) was added for one hour prior to the assay.

### C32 melanoma cell binding assay

C32 melanoma binding assays were performed as previously described ^72^. To prepare iRBCs for the cellular inhibition assay, mature-stage iRBCs (3D7 or FCR3) were prepared as for the receptor binding assays above. Plasma gel-enriched FCR3 iRBCs were overlaid onto confluent C32 cell monolayers in 6-well plastic dishes for 45 minutes in binding medium. After a 1-hour incubation, unbound iRBCs were removed by gentle washes with RPMI medium adjusted to pH 6.8 until only adhering parasites were observed. For each taurine concentration, ten independent 1 mm² microscopic fields were counted across three biological replicates. iRBC adhesion was quantified by counting bound iRBC per mm^2^ of confluent C32 cells in 10 random fields. Representative micrographs and raw data are provided in the supplementary materials (Fig. 4D, S4D, SM6).

### Flow cytometry assessment of *var19* surface expression

Control and supplementation of [0.64mM] taurine in FCR3 *var19*-expressing parasites iRBCs at mid/late trophozoites stages were resuspended in DPBS 1% BSA after VARIOMACS purification and counted. For each assay, 3 ×10^5^ iRBCs were washed in DPBS and incubated with 50µl of purified rabbit anti-*var19* antibodies ^54^ diluted 1:100 in PBS 1% BSA for 1-hour at room temperature. iRBCs were washed twice with DPBS and resuspended in 100µl of PE conjugated goat anti-rabbit antibody diluted 1: 100 in DPBS for 30 min at RT. iRBCs were washed twice in DPBS 1% BSA and resuspended in paraformaldehyde 4% in DPBS and kept at 4°C overnight in darkness. Cells were washed twice with DPBS and analyzed by flow cytometry using a BD LSR Fortessa flow cytometer. The results were analyzed using the FlowJo 10.0 software. Parasite nuclei were stained by TO-PRO-3 (1:10000 dilution). The results are expressed as the percentage of *var19* surface expression. A total of three biological replicates (n=3) were performed.

### Statistical analysis

For metabolomics MS analysis, GRmeta (https://github.com/kroemerlab/GRMeta) was used for the data shape QC correction (based on global pool for normalization between 3 total batches).

Fold change (FC) was determined when comparing each condition of each month to all other conditions in each month using the following code:

*#Fold Change averages*
*mm = mergedata$Data$AreaCorrLog2Cen2*
*mm = cbind(as.data.frame(mergedata$Grp2),mm)*
*mm = apply(mm[,-1], 2, function(x) tapply(x, mm$‘mergedata$Grp2’, mean)) %>% t()*
*FC = combn(1:ncol(mm),2,function(x) mm[,x[2]] - mm[,x[1]])*

*p* values were determined when comparing each condition of each month to all other conditions in each month using Kruskall wallis with BH adjustment with the following code:

*KruskalWallis$data[[mergedata$Annot$MetName[i]]]=as.data.frame(dunn.test(mergedata$ Data$AreaCorrLog2Cen2[,i],mergedata$Grp2, method="BH", kw=TRUE, label=TRUE,wrap=FALSE, table=TRUE, list=FALSE, rmc=FALSE, alpha=0.05, altp=FALSE)$P.adjusted)*

Metabolite enrichment and pathway analyses was performed using the Over Representation and Quantitative Enrichment tools on the MetaboAnalyst 6.0 server (https://www.metaboanalyst.ca/), integrated with the KEGG database (https://www.genome.jp/kegg/). Input data included a list of compound names (KEGG and HMDB IDs) for metabolites passing GC-MS quality control, categorized by their specific class. Pathway-based analysis was performed with at least two entries in the metabolite set library, which includes 80 metabolite sets based on KEGG human metabolic pathways (Dec 2023). Results were visualized as dot plots displaying key information on enrichment ratios, p-values, and log2 values for the respective metabolite sets.

To identify metabolites associated with infection status while accounting for temporal and repeated-measures dependencies, we employed generalized linear mixed-effects models (GLMMs) with a binomial logit link. For symptomatic malaria (S vs AS), the model included the scaled metabolite level as a fixed effect and sample month (October, November) as a random intercept. For asymptomatic infection (AS vs NI), the model included the metabolite level as a fixed effect, with sample month (December, January, February) and individual included as random intercepts. For chronic asymptomatic (AS) infections across the dry season, the model was extended to test for temporal effect modification. It included the fixed effects of the metabolite level, time (month, from December to May), and their interaction term, with a random intercept for individual.

All other statistical analyses were performed using R version 4.3.2 and GraphPad Prism version 9.1.0 (216) for Mac. To test for a normal distribution of the data, the Shapiro-Wilk normality test was used. To test for significance between two groups, a two-sided independent-samples t test was used.

## Supplementary Materials

Fig. S1 to S4

SM1 Information on samples from volunteers (excel file)

SM2 Metabolite alternative names (excel file)

SM3 Metabolomics MS raw data (excel file)

SM4 Pathway Enrichment Analysis (excel file)

SM5 Fold Change, p-values, and results from additional analyses to determine significantly increased or decreased metabolites depending on severity and seasonality (excel file)

SM6 iRBC adhesion counting raw data (excel file)

**Fig. S1:**
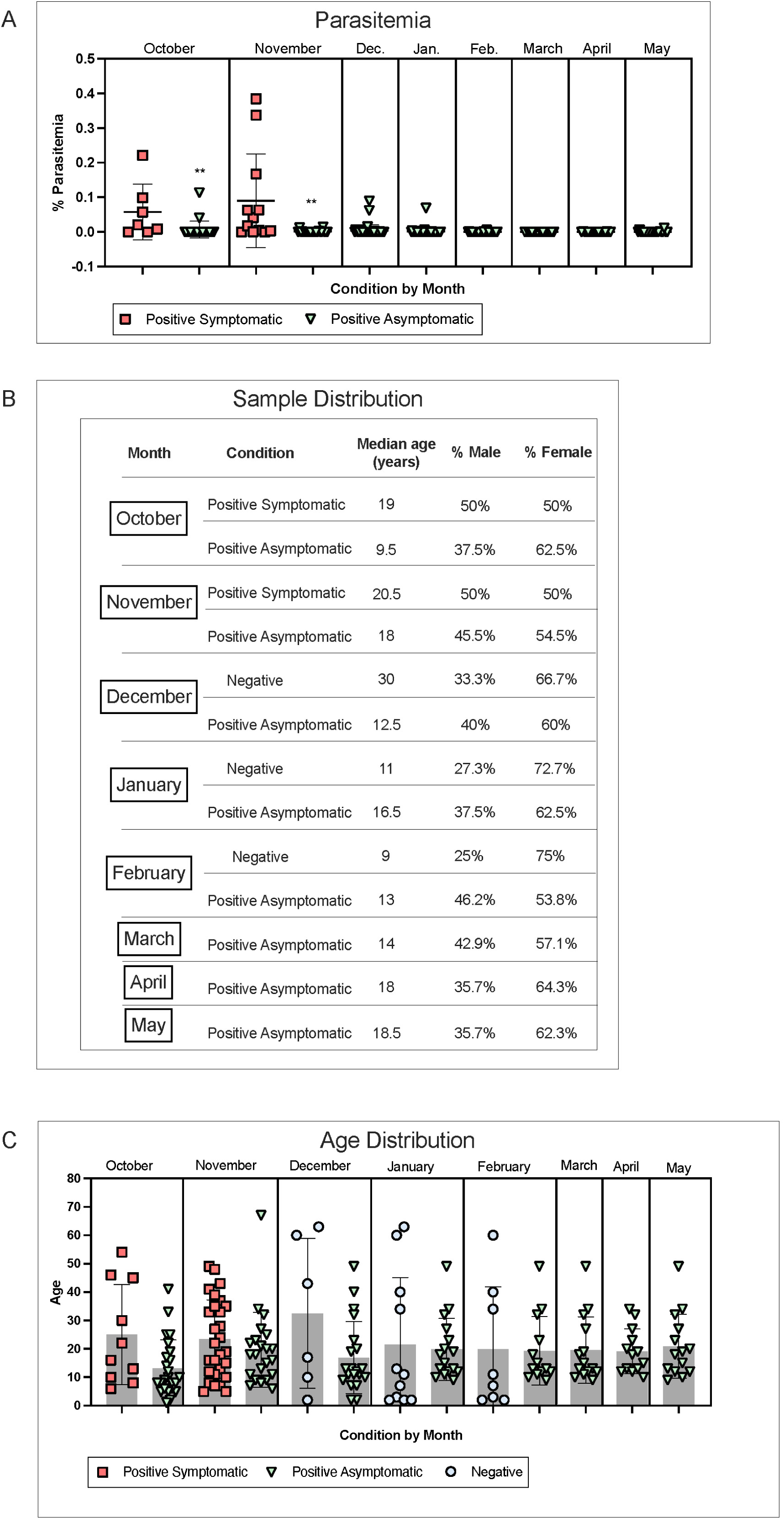
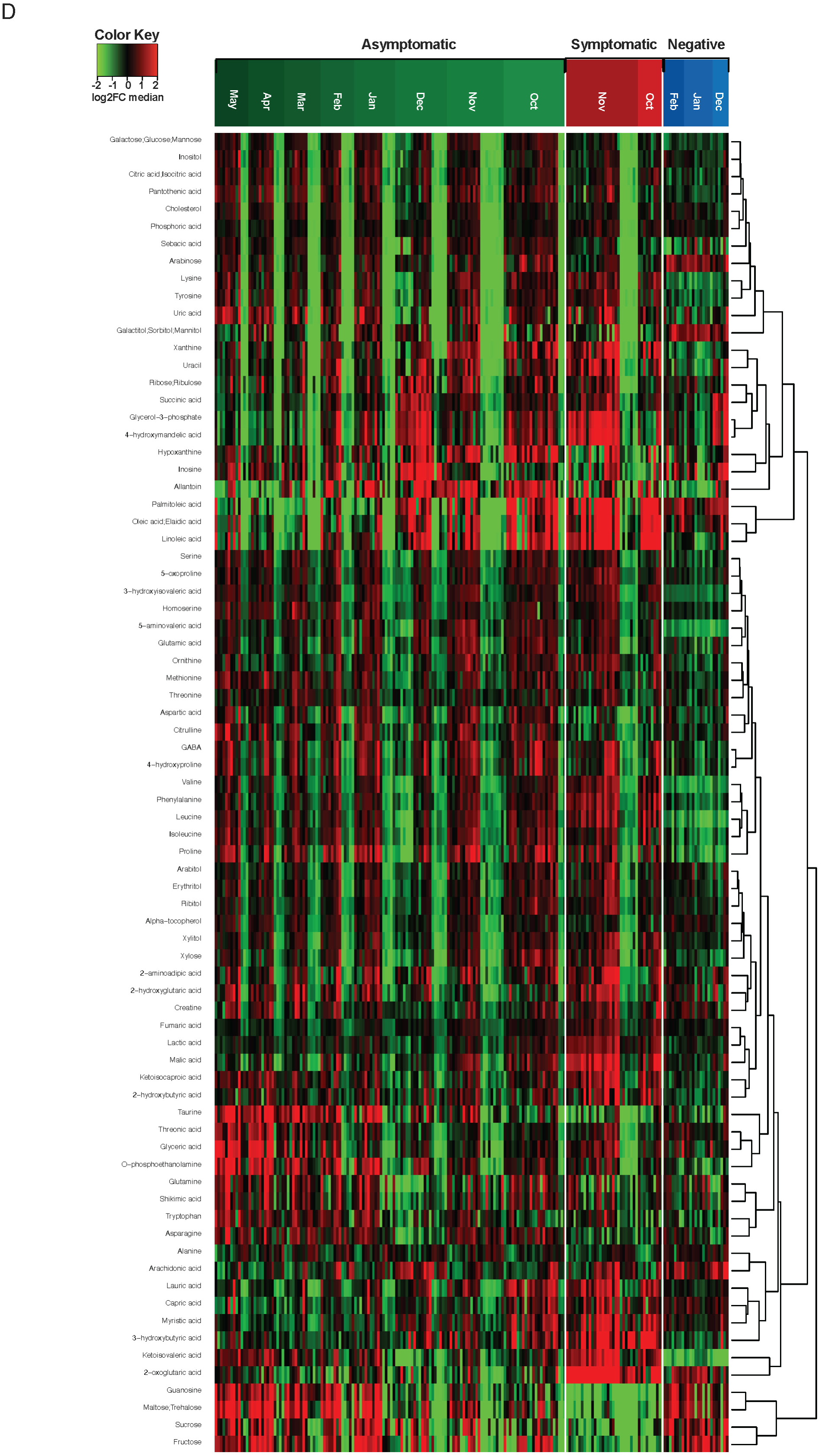
Additional information on samples used. (**A**) Parasitemia determined by qPCR for each of the conditions with available data, analyzed previously in ^44^. Symptomatic October (n=7) and November (n=12), Asymptomatic October (n=24), November (n=22), December (n=44), January (n=42), February (n=29), March (n=3), April (n=4), and May (n=21). The center line represents the mean with error bars showing standard deviation. Statistical significance was determined by unpaired t-test comparing symptomatic infections to asymptomatic infections in the same months (** October p = 0.0085, November p = 0.0038). (**B**) Sample distribution for all conditions in each month. The median age in years is shown as well as the distribution by sex for males and females. **(C)** Age distribution for all individuals from each of the conditions in each month. Grey boxes represent the mean age with error bars showing standard deviation. (**D**) Heatmap for all 78 metabolites and (n=199) plasma samples with hierarchical clustering illustrating the changes in metabolite abundances according to the median of each metabolite. Rows are metabolites, columns are samples. Heatmap data are log2 normalized and centered around the average abundance computed from all the samples for each metabolite. Red/green colors are ion signal higher/lower than average. Samples and metabolites are clustered following the ward.D2 algorithm, with Euclidean distance.

**Fig. S2:**
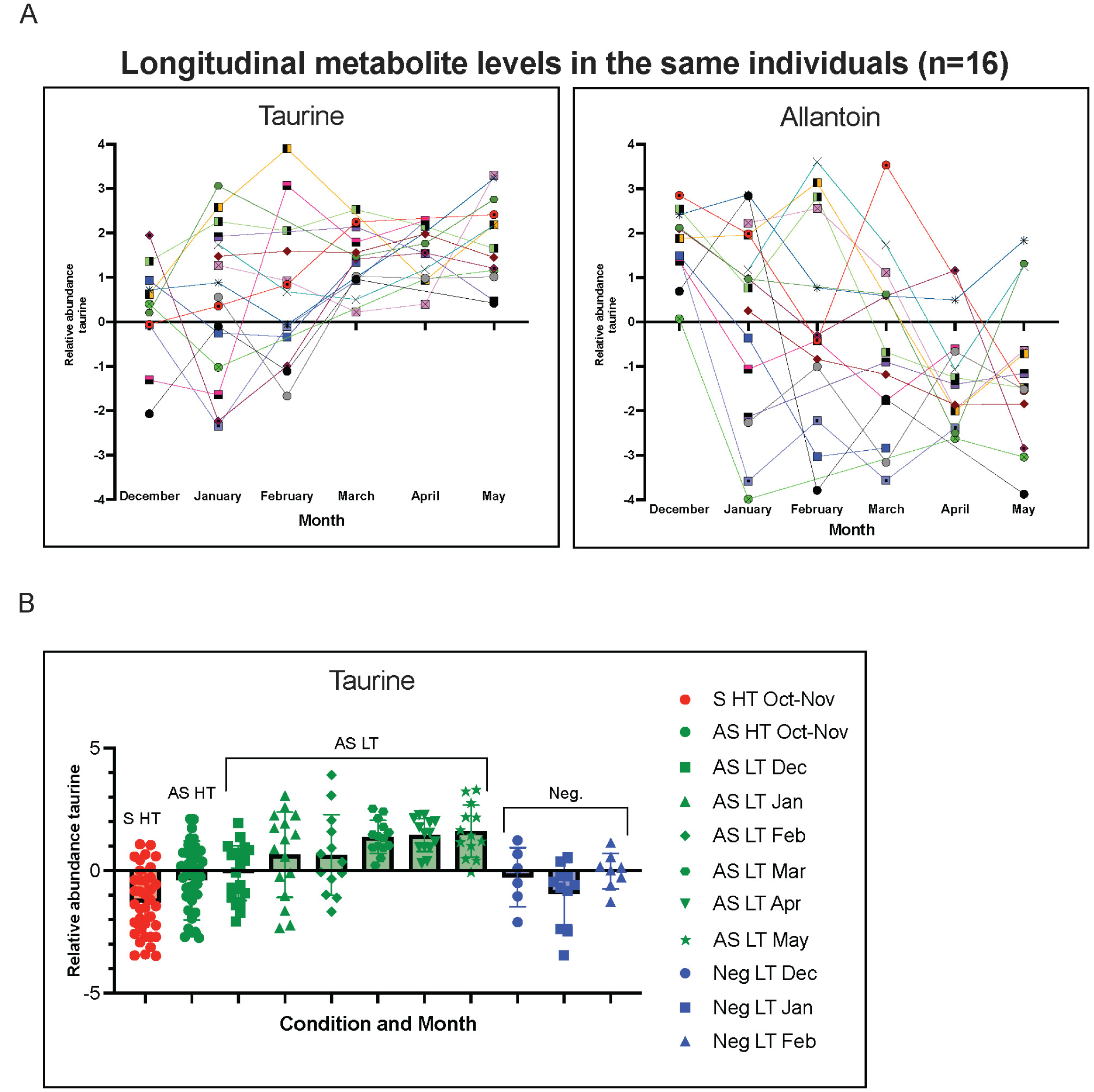
Taurine levels rise in multiple individuals during the low transmission period. **(A)** Spaghetti plots showing taurine and allantoin levels in individuals (n=16) with persistent infections who had samples taken for consecutive months during the low transmission period (December-May). Values plotted are relative abundance determined from the MS analysis. **(B)** Scatter plot showing relative abundance levels, determine by MS metabolomic analysis, of taurine in symptomatic infections, asymptomatic infections, and uninfected negative individuals. High transmission (HT) and low transmission (LT) months are labeled. Error bars represent standard deviation and mean values are represented in shaded boxes for all conditions and corresponding months.

**Fig. S3:**
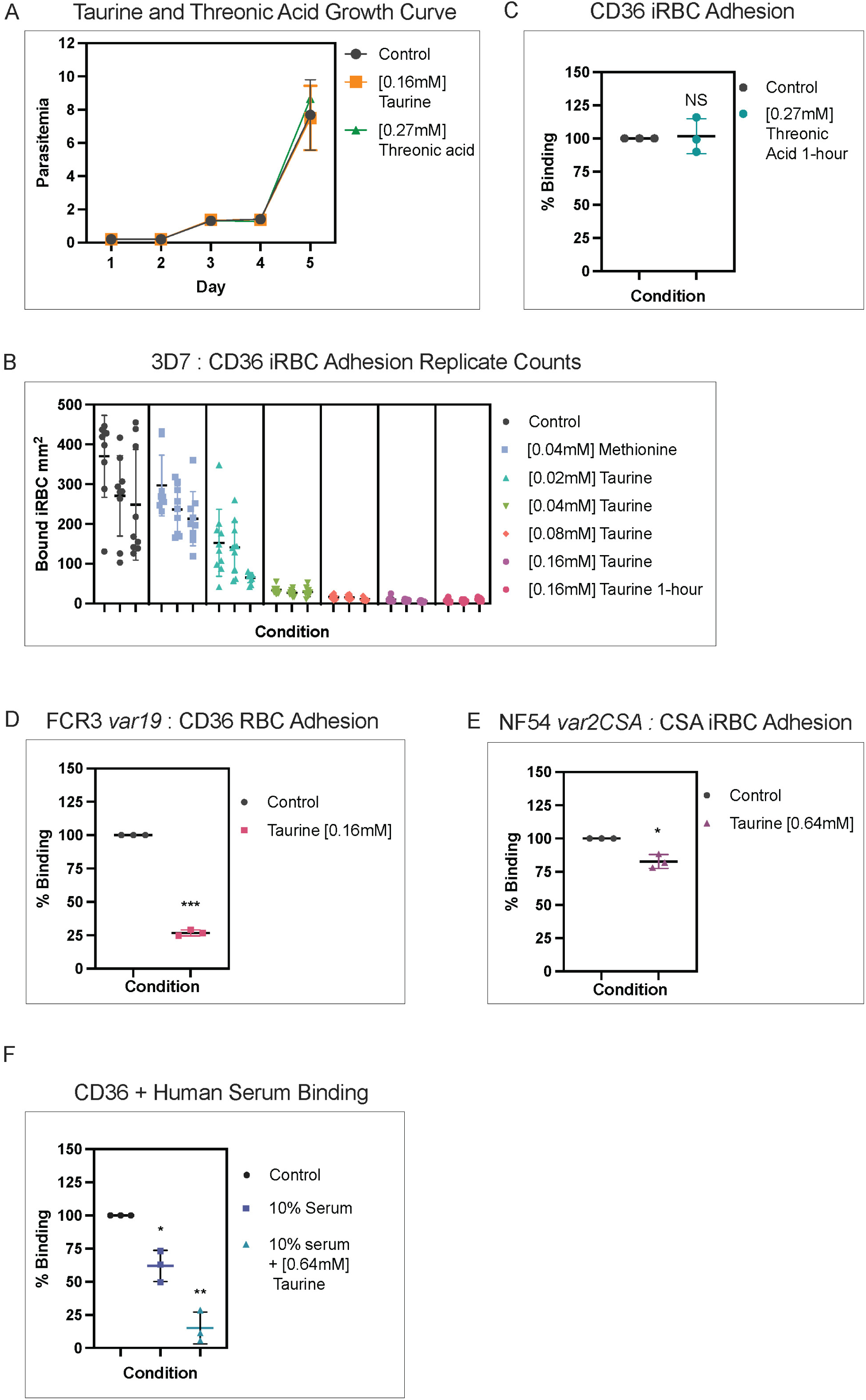
Taurine reduces iRBC adhesion without affecting parasite growth. (**A**) Five-day growth curve of 3D7 wild-type *P. falciparum* cultured under three conditions: control, 0.16 mM taurine (physiological concentration), and 0.27 mM threonic acid (physiological concentration). Error bars represent the standard deviation of three biological replicates performed in triplicate with different donor blood (n = 3). (**B)** Replicate counts (n=3) for adhesion assays from (Fig. 3A). Data is expressed as the number of bound 3D7 iRBCs per mm^2^ of coated receptor for each condition. **(C**) iRBC adhesion assays showing *P. falciparum* 3D7 iRBCs binding to CD36 after 1-hour incubation with 0.27 mM threonic acid. Data are presented as the percentage of iRBCs bound per mm² (±SD), relative to control; median values are indicated by the center line. Results represent three biological replicates (n = 3), each performed in duplicate. Statistical significance was determined by paired t-test (* p = 0.0305 and 0.0152, **** p < 0.0001). NS indicates not significant. **(D)** iRBC adhesion assays examining parasite FCR3 Itvar19 iRBCs binding to CD36, with or without pre-treatment with [0.16mM] taurine for one hour before the binding assay [parasites were the same strain as those used from (Fig. 3C)]. Data is expressed as the percentage of bound iRBCs ±SD per mm^2^ for each condition compared to control, with the median represented at the center line. Results are from a total of three biological replicates (n=3) done in duplicate. Statistical significance was determined by paired t-test (*** p = 0.0003). (**E)** iRBC adhesion assays for NF54 VAR2CSA iRBCS binding to CSA for control and [0.64mM] taurine. Data is expressed as the percentage of bound iRBCs ±SD per mm^2^ for each condition compared to control, with the median represented at the center line. Results are from a total of three biological replicates (n=3) done in duplicate. Statistical significance was determined by paired t-test (* p = 0.0288). **(F)** iRBC adhesion assays assessing 3D7 iRBCs binding to CD36 under control conditions and with the addition of 10% human serum with or without [0.64mM] taurine supplementation. Data is expressed as the percentage of bound iRBCs ±SD per mm^2^ compared to control, with the median represented at the center line. Results are from a total of three biological replicates (n=3) done in duplicate. Statistical significance was determined by paired t-test (* p= 0.035, ** p = 0.0066).

**Fig. S4:**
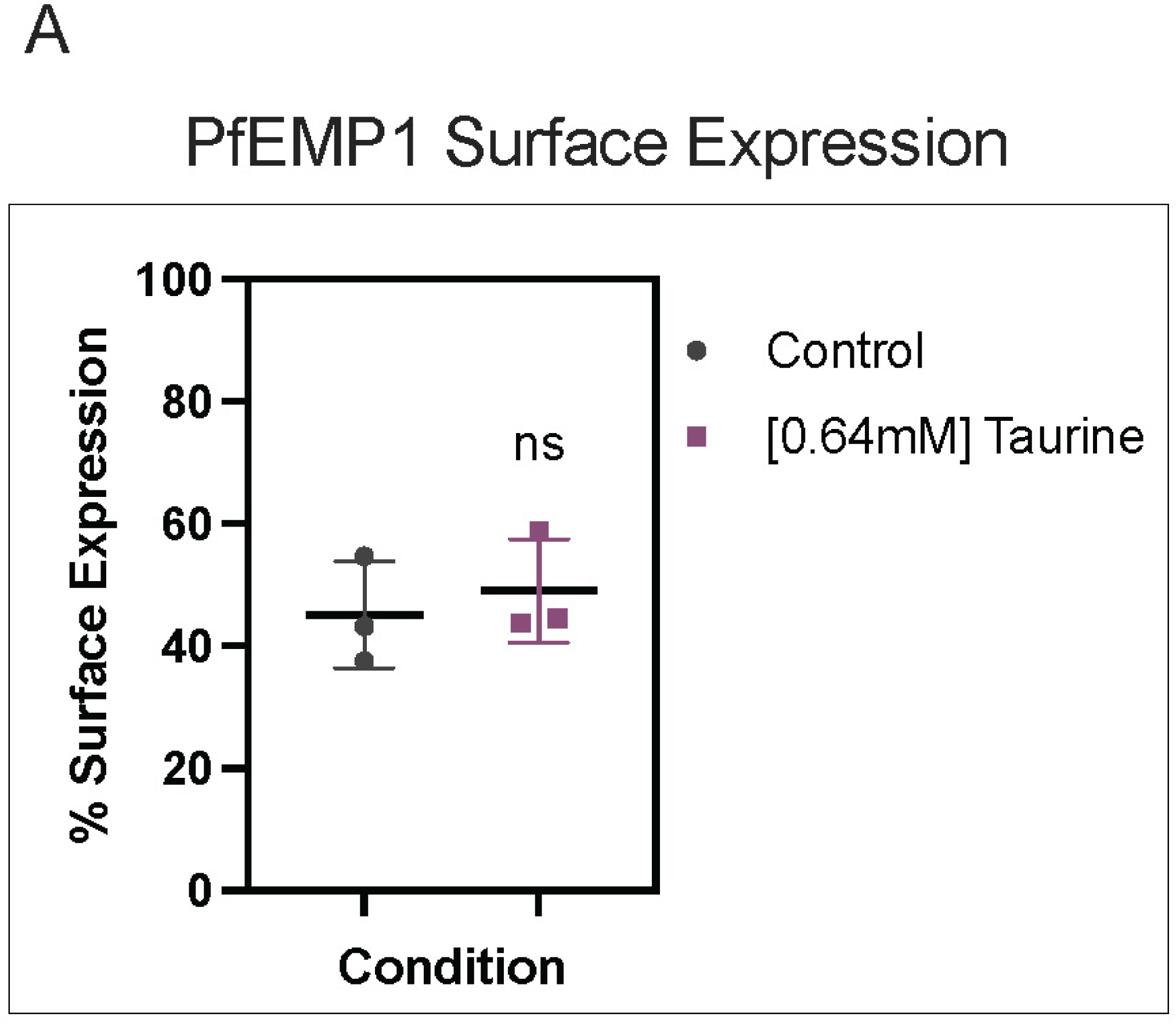
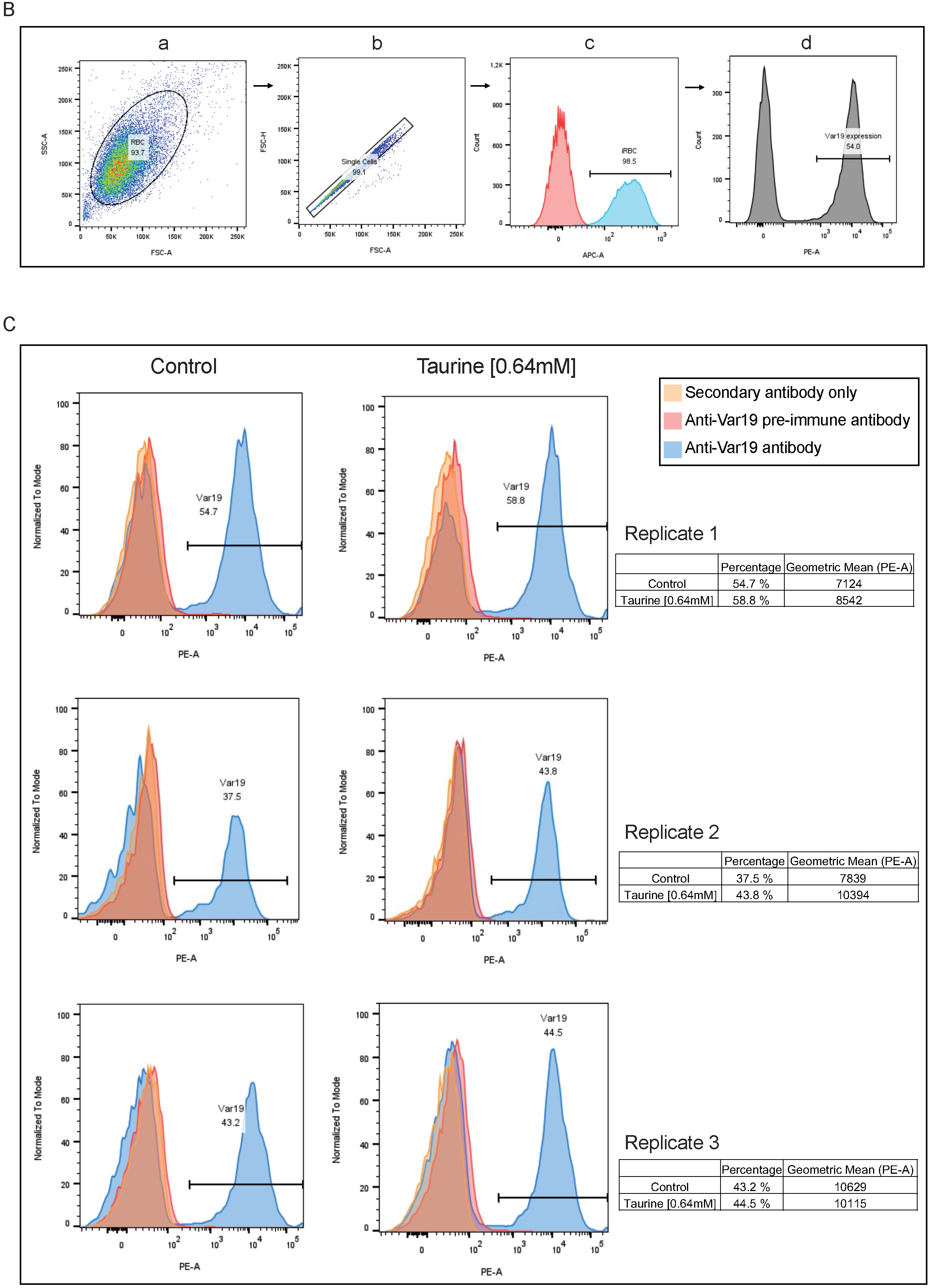
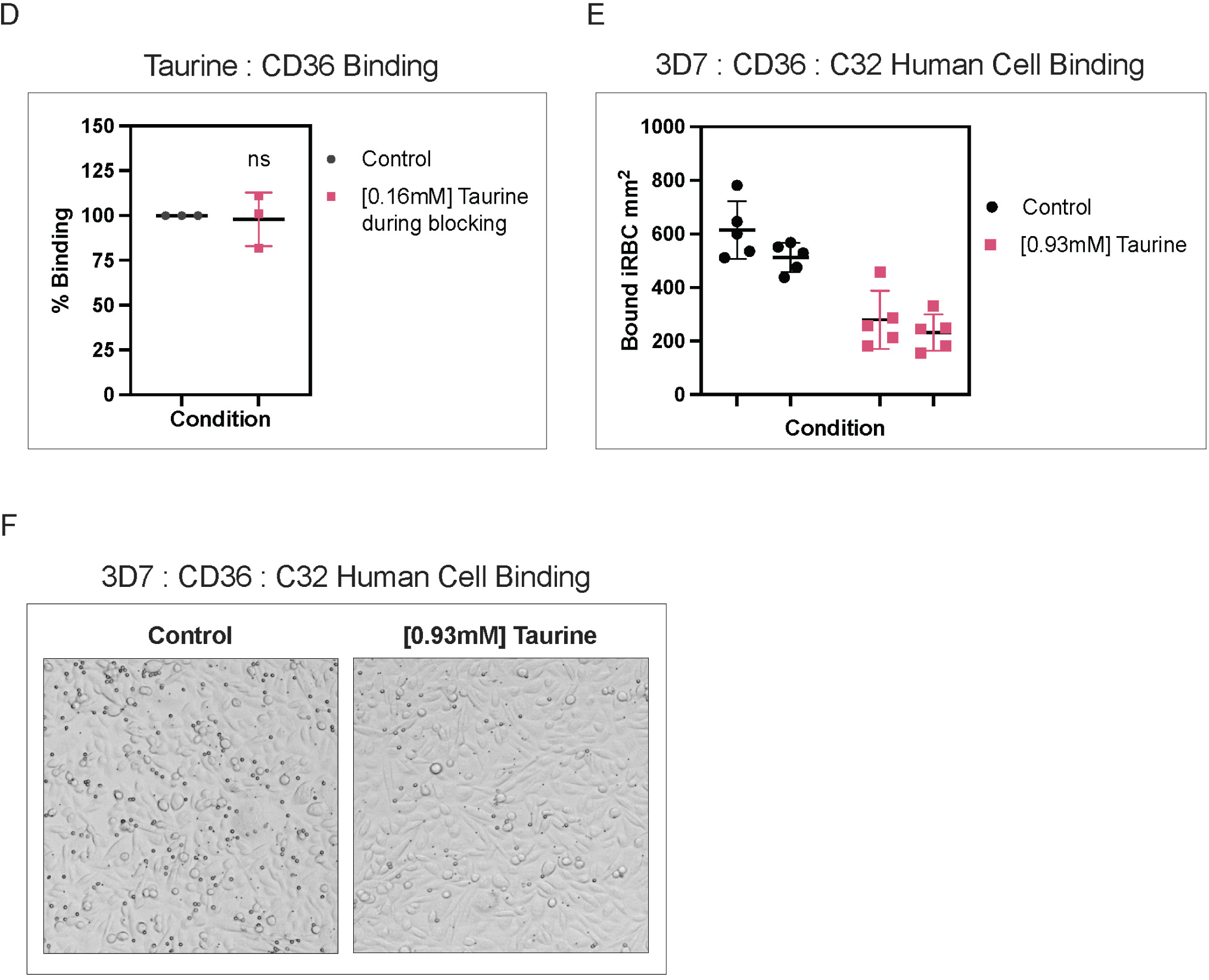
Taurine does not alter PfEMP1 surface expression. **(A)** FACS analysis for surface expression of PfEMP1 from parasites, incubated with or without [0.64mM] taurine, from a total of three biological replicates (n=3) done in duplicate using an antibody against *var19* PfEMP1. Statistical significance was determined by paired t-test (NS indicates not significant). (**B**) Flow-cytometry gating strategy. Gating hierarchy for a representative sample. From left to right: (a) In the FSC-A versus SSC-A plot, the cloud of RBC events is identified. (b) Among the RBCs, in the FSC-A versus FSC-H plot, singlets (single cell signals) are distinguished. (c) Among the singlets, the APC-A histogram is used to identify iRBCs. (d) Finally, in the PE-A channel, cells expressing var19 on their surface are identified based on PE-A labelling. **(C)** Flow cytometry analysis of FCR3 Var19 infected red blood cells. Percentage of Var19 expression on the cell surface without or with [0.64mM] taurine supplementation (negative controls included). Histogram legends for the expression data and the percentage of Var19 expression on the surface with the geometric mean are shown. Results are from three biological replicates (n=3). **(D)** iRBC adhesion assays assessing 3D7 parasite binding to CD36 under control conditions and with CD36 pre-incubated with [0.16mM] taurine. Data is expressed as the percentage of bound iRBCs ±SD per mm^2^ compared to control, with the median represented at the center line. Results are from a total of three biological replicates (n=3) done in duplicate. Statistical significance was determined by paired t-test (NS indicates not significant). **(E)** Cytoadhesion assays measuring 3D7 iRBCs binding to human C32 melanoma cells expressing CD36 and ICAM-1 under control conditions and [0.93mM] taurine. Replicate counts (n=2) for cytoadhesion assays. Data is expressed as the number of bound iRBCs per mm^2^ of confluent C32 cells for each condition (±SD), with medians indicated by center lines. (**F**) Representative field of cytoadhesion assay from (Fig. S4E).

## Acknowledgments

We thank Jessica Bryant for critical reading of this manuscript. We thank Célia Peuziat and Sooraj Shivakumar for technical assistance with this project.

## Funding

This work was supported by funding to AS from the Laboratoire d’Excellence (LabEx) ParaFrap (ANR-11-LABX-0024), ANR VAR2PTM (ANR-22-CE15-0023). This work was also supported by funding to AC from MRC/LSHTM fellowship, the Agence Nationale de la Recherche (ANR-18-CE15-0009-01). This work was additionally supported by funding to BG from the Fondation pour la Recherche Médicale (Equipe FRM EQU202203014741).

## Author contributions

Conceptualization: GD, AS

Methodology: GD, PM, PC

Investigation: GD, PM, PC, BF, FA, DD, SD

Visualization: GD

Funding acquisition: AS, AC, BG

Project administration: GD, AS

Supervision: AS, AC, BG

Writing – original draft: GD, AS

Writing – review & editing: GD, AS, AC, BG

## Competing interests

The funders had no role in the study design, data collection, and interpretation or decision to submit the work for publication. The authors declare that they have no conflict of interest.

## Data and materials availability

All data are available in the main text or the supplementary materials.

## Notes

### Competing Interest Statement

The authors have declared no competing interest.

### Summary of Updates

Changes have been made to the results section and corresponding figures.

